# The uneasy coexistence of predators and pathogens

**DOI:** 10.1101/721506

**Authors:** Andreas Eilersen, Kim Sneppen

## Abstract

Disease and predation are both highly important in ecology, and pathogens with multiple host species have turned out to be common. Nonetheless, the interplay between multi-host epidemics and predation has received relatively little attention. Here, we analyse a model of a predator-prey system with disease in both prey and predator populations and determine reasonable parameter values using allometric mass scaling relations. Our analysis focuses on the possibility of extinction events rather than the linear stability of the model equations. We find that if the predator is a specialist, epidemics frequently drive the predator species to extinction. If the predator has an additional, immune prey species, predators will usually survive. Coexistence of predator and disease is impossible in the single-prey model. We conclude that for the prey species, carrying a pathogen can be an effective weapon against predators, and that being a generalist is a major advantage for a predator.

## INTRODUCTION

Predation is one of the fundamental modes of interaction among living organisms. Mechanisms similar to predation are found in anything from mammals to bacteria. Another equally important factor is epidemic disease, which is also found on all scales in the ecosphere. In recent years it has become clear that many epidemic pathogens are shared between several species [1], of which some presumably prey on each other. If the predator runs a risk of becoming infected when eating infected prey, it is possible that the prey species will be able to use the pathogen as a weapon against the predator. This could even be a very effective evolutionary strategy, given that prey species are often much more numerous than their predators, leading to a high infection pressure against the predator species [2]. On this basis, we propose the hypothesis that a disease shared between a prey species and its predator will often turn out to be a major problem for the predator, and thus perhaps a long term advantage for the prey. However, if the predator has several prey options, epidemics should pose much less of a threat to it, as it can just feed on an immune prey species in the event of an epidemic.

The dynamics of predator-prey-pathogen interactions in general have received some attention in recent decades. Most attention has been given to the interaction between predators and single-host epidemics or parasitism [3–7]. On the other hand, despite the ubiquity of shared pathogens, the interplay between predation and multi-host disease has not been as thoroughly studied. Nonetheless, a few models similar to the one we will put forward in this paper do exist. Hsieh and Hsiao [8] have constructed one such model, and Han *et al*. [9] briefly cover another. These examples focus their analyses on the linear stability of the fixed points of their system, whereas we will focus on extinction events. We choose this focus, since a linear stability analysis will not reveal if a population is temporarily driven to such low densities that it would lead to extinction in the real world despite the stability of a fixed point.

Furthermore, when analysing epidemiological models, it is difficult but crucial to determine what parameter ranges are realistic. A discussion of this problem is often missing from more theoretical treatments [8, 9]. Therefore, we will here attempt to use the allometric mass scaling laws for many demographic and epidemiological quantities to estimate the range of parameters.

It has long been known that quantities such as reproduction rate and metabolic effect scale with animal mass to some quarter power [10]. Attempts have been made in ecology to use this to predict the behaviour of predator-prey systems [11–13]. More recently, it has been shown that disease recovery and death rates also scale with animal mass [14], which is useful in epidemiological modelling [15]. The parameterisation that we will use here will be based in part on our previous work on parameterising the Lotka-Volterra predator-prey equations [13]. The mass scaling relations are for the most part fairly general across different classes of animals. We will here be using the mass scaling relations valid for mammals. One could construct similar models for predation among other animals by mainly changing the constants of proportionality [10], and we would therefore expect our model to be relevant even for non-mammals. Only when looking at entirely different organisms such as bacteria do we need to be more careful, as the mechanisms that might be responsible for the scaling are different [16]. Nonethe-less, a similar scaling law for metabolic effect exists even for bacteria [17]

In summary, the questions that we will try to answer here will be whether an epidemic affecting a prey species can drive a predator species to extinction, and if so, for what parameter values this will be most likely. We also want to examine the effect of a predator being a generalist, i.e. having an alternative prey option that is not affected by the epidemic.

## THE MODEL

The first classic theory upon which we will base our study is the Lotka-Volterra predator-prey model

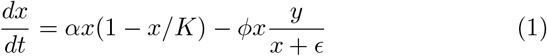

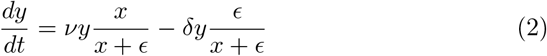

where *x* is prey, *y* is predator, *α* is the per capita prey reproduction rate and *δ* is the predator starvation rate in the absence of prey [18]. *K* is the prey carrying capacity and *ϵ* is the half-saturation constant for predators. The second term in the first equation is the functional response, which gives the rate of prey being eaten. We have here chosen to use a Holling type II functional response. Similarly, the first term of the second equation, 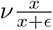, gives the predator reproduction rate as a function of prey population, the numerical response. We have modified the original Lotka-Volterra model to account for the natural limit on prey growth and the fact that predators do not grow infinitely fast when there is infinite available prey, nor do they starve when there is enough prey. We end up with a model very similar to the Rosenzweig-MacArthur model for predator-prey interactions [19].

We will combine these equations with the equally classic SIR model, which gives the following equations for the changes in population during an epidemic:

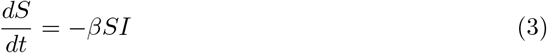

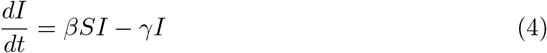

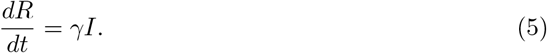

Here, *S* denotes susceptible individuals, *I* infected, and *R* recovered or dead individuals. *β* gives the rate at which each infected individual infects susceptible individuals, and *γ* gives the death or recovery rate of the infected [20].

When constructing our model, we shall make the assumption that the disease is always deadly, as the possibility of recovery with immunity will vastly complicate the analysis in a predator-prey system. Furthermore, we assume that infection from predator to prey is impossible, as any close encounters between the two species are likely to cause the immediate death of the prey. When modelling the system below, we find that varying the predator-predator infection rate makes relatively little difference. Figures illustrating this can be found in the supplement. For the sake of simplicity, in the following we will therefore only treat the case where the majority of predator infections stem from prey, and predator-predator infections can be neglected. We also let only healthy animals reproduce, although both healthy and infected predators eat prey. Combining the SIR and Lotka-Volterra models, we end up with the following equations for the single-prey system:

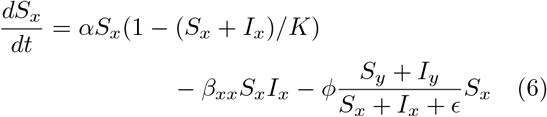

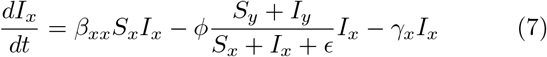

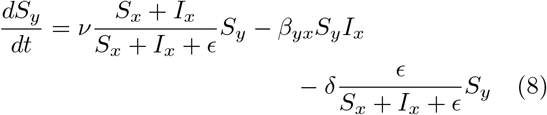

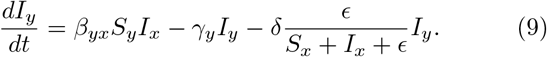

The equations for the number of dead individuals have been dropped, as they add no information when the disease is universally fatal. Subscripts here denote the species, with *β_ij_* being the coefficient for infection from species *j* to species *i*. If we set the probability of infection when eating an infected prey equal to 1, the infection coefficient *β_yx_* becomes equal to 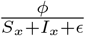, as the number of infected prey eaten equals the number of predators infected.

If we are to derive any information from these equations, it will be necessary to estimate realistic parameter values, and this is what we will do in the following.

From [10, 13] we can find relations between predator and prey mass (mx and my) and the parameters *α*, *δ*, *ϕ*, and *ν*. We want *α* and *ν* to represent theoretical maximal reproduction rates for prey and predators respectively. Instead of using the data from growing populations in the wild, where starvation, disease and other complications practically always play a role, we believe that the theoretical cap on reproduction should be set by the gestation period. *α* and *ν* should thus be the inverse gestation period [10]:

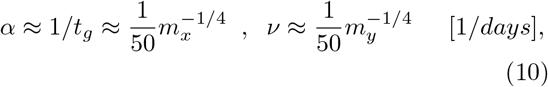

with mass in kilograms. A similar mass scaling law can be found for the incubation period of species that lay eggs [21]. Our intuition tells us that when the predator is satisfied (*S_y_* ≈ *ϵ*), predator reproduction should be equal to predator death, giving us 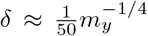 as well. In order to calculate how many prey the predators need to eat to reproduce this much, we must know the ecological efficiency *η*. The ecological efficiency, defined as the fraction of consumed prey biomass converted into predator biomass, we estimate to be 10 % although the quantity varies significantly with trophic level and the specifics of the species [22, 23]. Knowing the efficiency, we can calculate the number of prey eaten as 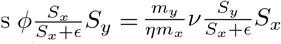, which implies 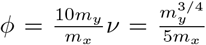 Finally, also from Peters [10], we have the following approximate relation for the carrying capacity:

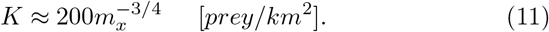

This relation is valid if we assume that the prey is a mammal and accept that the metabolic scaling exponent 3/4 is the “true” theoretical value of the empirically estimated scaling exponent (∼ 0.61) of the carrying capacity. By using this carrying capacity relation, we decide that the units of the population densities are [*km*^-2^]. *ϵ* is difficult to determine, and we therefore choose to set *ϵ* = *K*/2. We believe this to be reasonable, as it allows the predator population growth to saturate before the prey population reaches its carrying capacity. However, as can be seen in the supplement, we can set *ϵ* to practically any value between 0.3*K* and *K* and still get similar results.

To extend the predator-prey model to the predatorprey-disease case, we also need to know the scaling relations for disease duration. According to Cable *et al*. [14], both the time until first symptoms and the time until recovery or death scale as *t* = *cm*^1/4^, where *c* is an experimental constant. Here, we shall use the constants appropriate for rabies. We choose to use these constants since we need an estimate of the order of magnitude of the scaling coefficient for the infective period, and rabies behaves a lot like we assume the disease to do in our model (universally fatal, multiple host species, etc. [24]). It should be stressed, however, that the disease is not in fact rabies, since we also wish to study the effects of varying the infectivity of the disease.

According to Cable *et al*. the duration of the period during which the infected individual is symptomatic can be written *t_I_* ≈ *t_D_* − *t_S_* = (*c*_2_ − *c*_1_)*m*^1/4^, where *c*_1_ and *c*_2_ are the scaling coefficients appropriate for the time until first symptoms and death, respectively. We assume that this period is of the same order as the infective period of the disease. The constants have been determined using statistical analysis, and their values are *c*_1_ = 9 (4, 19) and *c*_2_ = 16 (7, 32), where the numbers in parentheses are the boundaries of the confidence interval from *p* = 2.5 % to *p* = 97.5 % [14, 24]. *γ_i_* can now be found as 1/*t_I,i_*.

Finally, to make the parameterisation more intuitive, we choose to express infectivity in terms of a quantity *R_xx_* related to the basic reproduction number (*R*_0_) of the disease. The basic reproduction number represents the number of secondary infections that occur when exposing an infected individual to a completely susceptible population. The reproduction number is related to the infection coefficient as 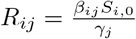 in the SIR model [2], where *S*_*i*,0_ is the initial density of susceptible individuals of species *i* at the onset of the epidemic. *R*_0_ ranges from 1, where an epidemic is barely able to sustain itself, up to 18 in measles [25]. We therefore intend to vary *R_xx_* from 1 to 10, as this covers a wide range of all possible disease infectivities. The cross-species reproduction number *R_yx_* will be determined by the number of prey eaten by predators which in turn depends on their mass ratio. Since our model is quite different from the SIR model, we should argue why the parameter *R_xx_* is equivalent to the original *R*_0_. *R*_0_ has the important property that if it is less than 1, the epidemic always dies out. If we choose the starting population *S*_*x*,0_ to be the Lotka-Volterra equilibrium in the absence of disease, we have 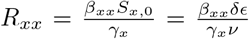. If we analyse the linear stability of the fixed point with coexsistence of prey and disease in the model with no cross-species infection, we find that this fixed point indeed becomes stable when *R_xx_* becomes greater than 1. The disease is thus able to persist only when *R_xx_* > 1. This means that *R_xx_* behaves like *R*_0_, and it is justified to use *R_xx_* as a basic reproduction number. As the initial predator population, we similarly choose the LV-equilibrium value, 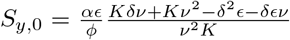, which reduces to 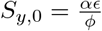 given our parameterisation.

By using this parameterisation, we are now left with only five parameters: Prey mass, predator mass, *ϵ*, prey-prey disease reproduction number, and the infection probability when predators eat infected prey. If we fix this probability at 1, we save another parameter. This is not always a good approximation [26]. However, varying the infection probability has a much smaller effect than varying *m_i_* or *R_xx_*, as is demonstrated in the supplement. We therefore choose to fix the probability at 1. Figures demonstrating this are shown in the supplement. The mass parameterisation further ensures that the values of the parameters used are at least biologically plausible. All this will be highly advantageous when we examine parameter space.

## EXAMINING PARAMETER SPACE

To derive information about parameter space most effectively, we perform a parameter sweep where we let the different quantities vary logarithmically. We scan a region of parameter space large enough that the species falling within this region are interestingly different. An overview of the parameter ranges and increments can be seen in table I. As initial condition, we choose the classical Lotka-Volterra equilibrium to avoid introducing large artificial oscillations into the system.

**TABLE I.** (a) The ranges of parameters examined in our logarithmic parameter sweep. *m_i_* is the mass of species *i*, and *R_xx_* is the disease reproduction number characterising infection from prey to prey. 330 parameter sets were tested.

|  | Min. value | Increment | Max. value |
| --- | --- | --- | --- |
| $\log(m_x)$ | -2 | 0.8 | 2 |
| $\log(m_y)$ | -1.2 | 0.8 | 2 |
| $\log(R_{xx})$ | 0 | 0.1 | 1 |

In the analysis of our model, we have found that when using reasonable parameter values, the populations will usually perform damped oscillations of initially large amplitude around some equilibrium, or approach a limit cycle with some possibly large amplitude. Although the equilibria might be stable, after a disease outbreak the populations often temporarily reach such low values that it would lead to extinction in any system with a discrete number of individuals. Also, an equilibrium with a very low population density would lead to extinction in the real world. If we introduce an extinction threshhold to our model to account for the finite and discrete number of individuals found in real populations, this changes the dynamics significantly.

We introduce an extinction threshhold of 10^−5^. If a population dips below this value, we consider it extinct. It should be noted that the precise value of the thresh-hold makes a relatively small difference in the end result. After solving the equations numerically over *T* = 20000 days, we classify the end state of the system into one of four categories: Scenarios with predator survival, disease persistence, disease-predator coexistence, and scenarios where only the healthy prey population survives. Since coexistence of disease and predator appears to be transient, we let the simulation run up to 10^5^ days if there is still coexistence at the end of the first simulation. Plots of the regions of parameter space with predator survival and disease persistence can be seen in fig. 1.

**FIG. 1.**
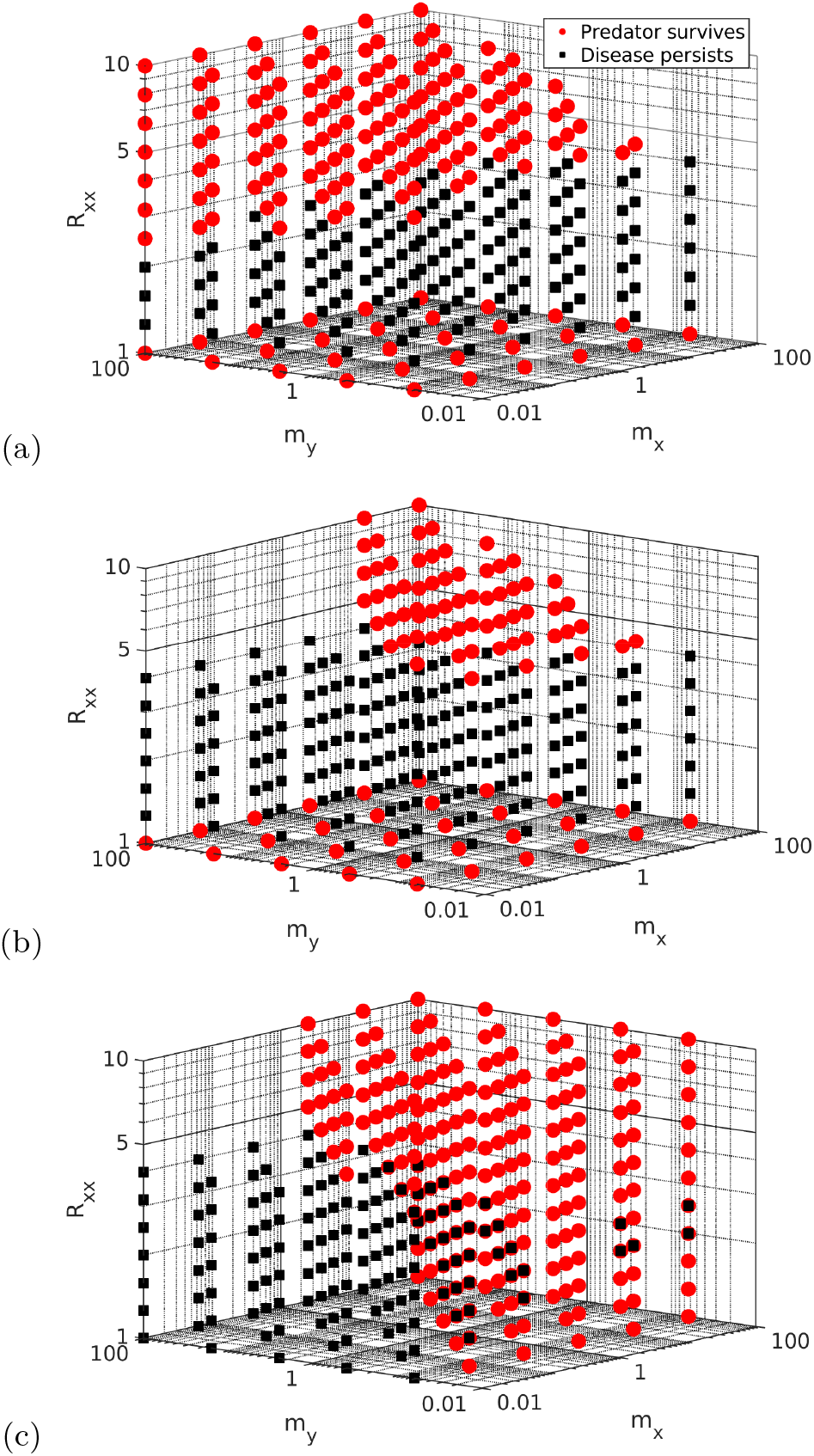
(Colour online) Parameter space regions where the predator survives or the disease persists, as a function of prey mass *m_x_*, predator mass *m_y_* (both in kg), and disease reproduction number between prey *R_xx_*. The coordinates of each red dot indicates a set of parameter values where the predator survives, while the location of the black squares indicate parameter values with disease persistence. If (a) the predators are immune, large predators survive at high *R_xx_*. On the other hand, if (b) the predators are susceptible, they survive if they are relatively large and subsist on a prey that is not too small. In (c), an immune prey is included alongside the susceptible one. The susceptible predators survive regardless of disease infectivity, so long as the predator is not too large compared to the prey. If the predator is not susceptible, it always survives (figure not shown).

First and foremost, we observe that the prey species practically always survives the epidemic. Therefore, there are a few scenarios where the prey ends up with no natural enemies and thus will grow to the carrying capacity and dominate the ecosystem. As another point, it should be noted that the pathogen never coexists with the specialist predator.

From the plots, we see that the survival of predators is strongly dependent on the reproduction number of the disease among prey and the mass of the predator species. When the epidemic does not directly affect the predators (fig. 1 (a)), the predators can survive at high *R_xx_*. A high mass is advantageous to the predator, as this ensures that it is not affected as much by starvation when the epidemic lowers the prey population. We see a “zone of exclusion” at intermediate *R_xx_* where the disease keeps the predator away. This gap is evidence of competitive exclusion between predator and disease, which both subsist on the same resource, the susceptible prey.

In the case where predators are susceptible to the disease (fig. 1 (b)), on the other hand, the predator species can sometimes survive at high *R_xx_*, but only if its mass is not too large. The predators survive if they have less than about 10 times the prey mass. Above this threshhold, the predator needs to eat so many prey to survive that it is almost guaranteed to be infected. These diagrams show us that sharing a pathogen with a prey species will most often cause the predator to go extinct. In fact, even an outbreak of a prey-specific epidemic can cause predator extinction, at least if the predator is a specialist.

The effect of additional prey species could be interesting to study, since a large part of the reason why epidemics are so deadly to predators appears to be that specialist predators either starve if prey population drops, or have to eat infected prey to avoid this. We do not expect these effects to be present, or at least not nearly as powerful in the presence of two prey species, of which one is immune. To test this hypothesis, we modify the equations (6)-(9) to include another prey that is unaffected by the disease. We assume that the immune prey is similar to the susceptible prey and simply set their parameters to be equal. The initial combined prey population is the same as before, and the prey species do not compete. We get the results seen in fig. 1 (c)

There is a striking difference compared to the case with only susceptible prey. In the two-prey model, predators will always survive, so long as they are small enough (less than 50-100 times the prey size at high *R_xx_* and less than the prey size at low *R_xx_*). We here see the same effect as in fig. 1 (b), that predators bigger than this need to eat a lot of prey and will therefore almost always end up eating an infected individual. If we instead assume that predators are immune, they always survive.

## DISCUSSION

The most striking conclusion to be drawn from this study is that an emerging epidemic in a specialist predator-prey system will tend to drive the predator, *but not the prey*, to extinction. Packer *et al*. have previously concluded that there are many situations in which a predator species might keep prey epidemics and parasites in check [7]. The argument that we will make based on this study is the converse: Given our dynamical model and an extinction threshhold, epidemic pathogens will make life hard for predators. The parameter sweeps show that disease and specialist predators cannot coexist. We believe this to be an example of competitive exclusion. Both the pathogen and the predator share a resource - the susceptible prey - and in such cases, longterm coexistence is impossible [27]. As the spread of the disease is not limited by saturation or energetic concerns, it will tend to win over the predator. The obvious implication of these two conclusions is that we should see very few ecosystems with specialist predators, prey, and a shared pathogen in the real world, as they are inherently unstable. Indeed, when examining existing literature, we have found no examples of specialist predators and prey sharing a deadly pathogen, nor of less specialised predators sharing a pathogen with their main prey. Of course, we should be careful drawing too many conclusions from our inability to find such systems. For example, pathogens that do not affect humans or economically important species are likely to receive less attention and therefore feature less prominently in the literature. However, our model can be used to argue that predatorprey-pathogen ecosystems should be rare, and we have so far not been able to falsify this. One further potential caveat is that we have here focused on mammals, using the mass scaling relations and assumptions relevant for mammalian predator-prey systems. However, due to the near-universality of mass-scaling relations in animals [17] we expect that most of the relations derived here should be easily transferable to other classes.

Quite often when the predator goes extinct but the epidemic persists, the resulting prey-pathogen equilibrium will have a lower prey population than the prey-predator equilibrium. Nonetheless, carrying the pathogen may still turn into an advantage for the prey species. From evolutionary biology, we know that when a pathogen becomes endemic in a given species, there will be a pressure for it to evolve to become less lethal over time [28]. This allows the pathogen to live longer in each host, and possibly to spread more effectively. An initially fatal epidemic can thus end up becoming harmless to its primary host species. If it has wiped out the predator in the process, this will represent a win-win situation for the prey species.

Possibly, we could expand our conclusion and simply state that specialist predators generally are extremely vulnerable to any changes in prey population. It is a well-known fact of ecology that specialists are the most vulnerable to extinction in the event of changes such as disease outbreaks [29]. Predator species that subsist entirely on one prey species should therefore also be rare in nature. This is supported by the fact that even predators which are usually noted as specialists, such as the weasels (*Mustela nivalis*) of northern Scandinavia that often form the basis of predator-prey models [30], tend to switch prey in times when their preferred prey is scarce [31].

Finally, as an additional result, predators that are much bigger than the size of their prey are a lot more vulnerable to infection with a shared pathogen from their prey, since they need to eat more potentially infected individuals to survive. This is true even for generalist predators and is an obvious consequence if a large percentage of the prey population is infected. What is less obvious is that the upper bound on predator to prey mass ratio drops abruptly when *R_xx_* dips below a certain threshhold (around 4) in the generalist predator case. Above this threshhold, a generalist predator species can be nearly a hundred times the size of its infectious prey species and still not go extinct due to infection. Below the threshhold, any predator larger than the infected prey will be driven to extinction by cross-species infections. The reason behind this change is that at high infectivities, the epidemic quickly uses up the supply of susceptibles and dies out. Therefore, the entire predator population will not have time to be infected. This result, in addition to energetic concerns about hunting very small animals, could lead to an evolutionary pressure for predators to not grow too large compared to their prey.

Given all of the above, we conclude that epidemic diseases can serve as an evolutionary weapon against specialist predators. A pathogen infecting a prey species will competitively exclude any specialist predator species, even when the predator is not itself susceptible to the pathogen. Shared epidemics between predator and prey may help impose an upper limit on the predator-prey size ratio, since eating a lot of small prey is dangerous if the prey is infectious. The negative effect of prey disease on the predator is however weakened a lot when we take into account additional, immune prey species. The uneasy coexistence of predators and pathogens should make specialist predator-prey-disease systems rare in the real world, and this is apparently the case. Our study supports the conclusion that being a specialist predator is a highly vulnerable position, and that being a generalist should be evolutionarily favourable for predator species. Normally, one would expect that competitive exclusion presents a drive towards speciation and specialisation [32]. Our model, on the contrary, provides an example of how the inherent vulnerability of specialists will drive species towards generalisation.

In conclusion, our study supports the idea that shared epidemic diseases could be a much more important factor in the coevolution of predator and prey species than they are usually given credit for.

## Supporting information

Supplemental figures - values of epsilon

## ACKNOWLEDGEMENTS

Our research has received funding from the European Research Council (ERC) under the European Union’s Horizon 2020 research and innovation programme under grant agreement No. [740704].

## References

[1] Mark E. J. Woolhouse and Sonya Gowtage-Sequeria, “Host range and emerging and reemerging pathogens.” Emerging infectious diseases 11, 1842–1847(2005).

[2] Sergei Maslov and Kim Sneppen, “Severe population collapses and species extinctions in multi-host epidemic dynamics,” Phys. Rev. E 96 (2017).

[3] J. Chattopadhyay and O. Arino, “Predator-prey model with disease in the prey,” Nonlinear Analysis, Theory, Methods and Applications 36, 747–766(1999).

[4] Herbert W. Hethcote, Wendi Wang, Litao Han, and Zhien Ma, “A predator-prey model with infected prey.” Theoretical population biology 66, 259–268(2004).

[5] Pierre Auger, Rachid Mchich, Tanmay Chowdhury, Gauthier Sallet, Maurice Tchuente, and Joydev Chattopadhyay, “Effects of a disease affecting a predator on the dynamics of a predator-prey system.” Journal of theoretical biology 258, 344–351(2009).

[6] H. I. Freedman, “A model of predator-prey dynamics as modified by the action of a parasite.” Mathematical biosciences 99, 143–155(1990).

[7] Craig Packer, Robert D Holt, Peter J Hudson, Kevin D Lafferty, and Andrew P Dobson, “Keeping the herds healthy and alert: implications of predator control for infectious disease,” Ecology Letters 6, 797–802(2003).

[8] Ying-Hen Hsieh and Chin-Kuei Hsiao, “Predator-prey model with disease infection in both populations,” Mathematical Medicine and Biology: A Journal of the IMA 25, 247–266(2008).

[9] Litao Han, Zhien Ma, and H. W. Hethcote, “Four predator prey models with infectious diseases,” Mathematical and Computer Modelling 34, 849–858(2001).

[10] Robert Henry Peters, The ecological implications of body size (Cambridge University Press, Cambridge, 1983).

[11] P. Yodzis and S. Innes, “Body size and consumer-resource dynamics,” The American Naturalist 139, 1151–1175(1992).

[12] Joshua Weitz and Simon Levin, “Letter: Size and scaling of predator-prey dynamics,” Ecology Letters 9, 548–557(2006).

[13] Andreas Eilersen and Kim Sneppen, “Applying allometric scaling to predator-prey systems,” Phys. Rev. E 99 (2019).

[14] Jessica M. Cable, Brian J. Enquist, and Melanie E. Moses, “The allometry of host-pathogen interactions (allometry and disease),” PLoS ONE 2 (2007).

[15] A. Dobson, “Population dynamics of pathogens with multiple host species,” The American Naturalist 164, 64–78(2004).

[16] Geoffrey B West, James H Brown, and Brian J Enquist, “A general model for the origin of allometric scaling laws in biology,” Science 276, 122–126(1997).

[17] Lev Ginzburg, Mark Colyvan, et al., Ecological orbits: how planets move and populations grow (Oxford University Press on Demand, 2004).

[18] Alfred J. Lotka, “Analytical note on certain rhythmic relations in organic systems,” Proceedings of the National Academy of Sciences of the United States of America 6, 410–415(1920).

[19] Michael L Rosenzweig and Robert H MacArthur, “Graphical representation and stability conditions of predator-prey interactions,” The American Naturalist 97, 209–223(1963).

[20] William Ogilvy Kermack, A. G. McKendrick, and Gilbert Thomas Walker, “Contributions to the mathematical theory of epidemics. iii. further studies of the problem of endemicity,” Proceedings of the Royal Society of London A 141, 94–122(1933).

[21] James F Gillooly, Eric L Charnov, Geoffrey B West, Van M Savage, and James H Brown, “Effects of size and temperature on developmental time,” Nature 417, 70 (2002).

[22] Raymond Lindeman, “The trophic-dynamic aspect of ecology,” Ecology 23 (1942).

[23] P. A. Colinvaux and B. D. Barnett, “Lindeman and the ecological efficiency of wolves,” The American Naturalist 114, 707–718(1979).

[24] Alan C. Jackson, Research advances in rabies, 1st ed., Advances in virus research v. 79 (Elsevier/Academic Press, Amsterdam Boston, 2011).

[25] William J. Moss and Diane E. Griffin, “Global measles elimination,” Nature Reviews Microbiology 4 (2006).

[26] R. O. Ramsden and D. H. Johnston, “Studies on the oral infectivity of rabies virus in carnivora.” Journal of wildlife diseases 11, 318–324(1975).

[27] G. F. Gause, “Experimental analysis of vito volterra’s mathematical theory of the struggle for existence,” Science 79, 16–17(1934).

[28] P.W. Ewald, “Guarding against the most dangerous emerging pathogens: Insights from evolutionary biology,” Emerging Infectious Diseases 2, 245–257(1996).

[29] Michael Begon, John Lander Harper, and Colin R Townsend, Essentials of ecology, 2nd ed. (Blackwell Publishing, 2003).

[30] I. Hanski, L. Hansson, and H. Henttonen, “Specialist predators, generalist predators, and the microtine rodent cycle,” The Journal of Animal Ecology 60 (1991).

[31] Janne Sundell and Hannu Ylnen, “Specialist predator in a multispecies prey community: boreal voles and weasels,” Integrative Zoology 3, 51–63(2008).

[32] Robert Macarthur and Richard Levins, “The limiting similarity, convergence, and divergence of coexisting species,” The American Naturalist 101, 377–385(1967).

