## Supplemental figures - values of epsilon for "The uneasy coexistence of predators and pathogens"

### Supplemental figures to the article ”The uneasy coexistence of predators and pathogens”

February 5, 2020

### Outcomes for different values of the half-saturation constant $\epsilon$

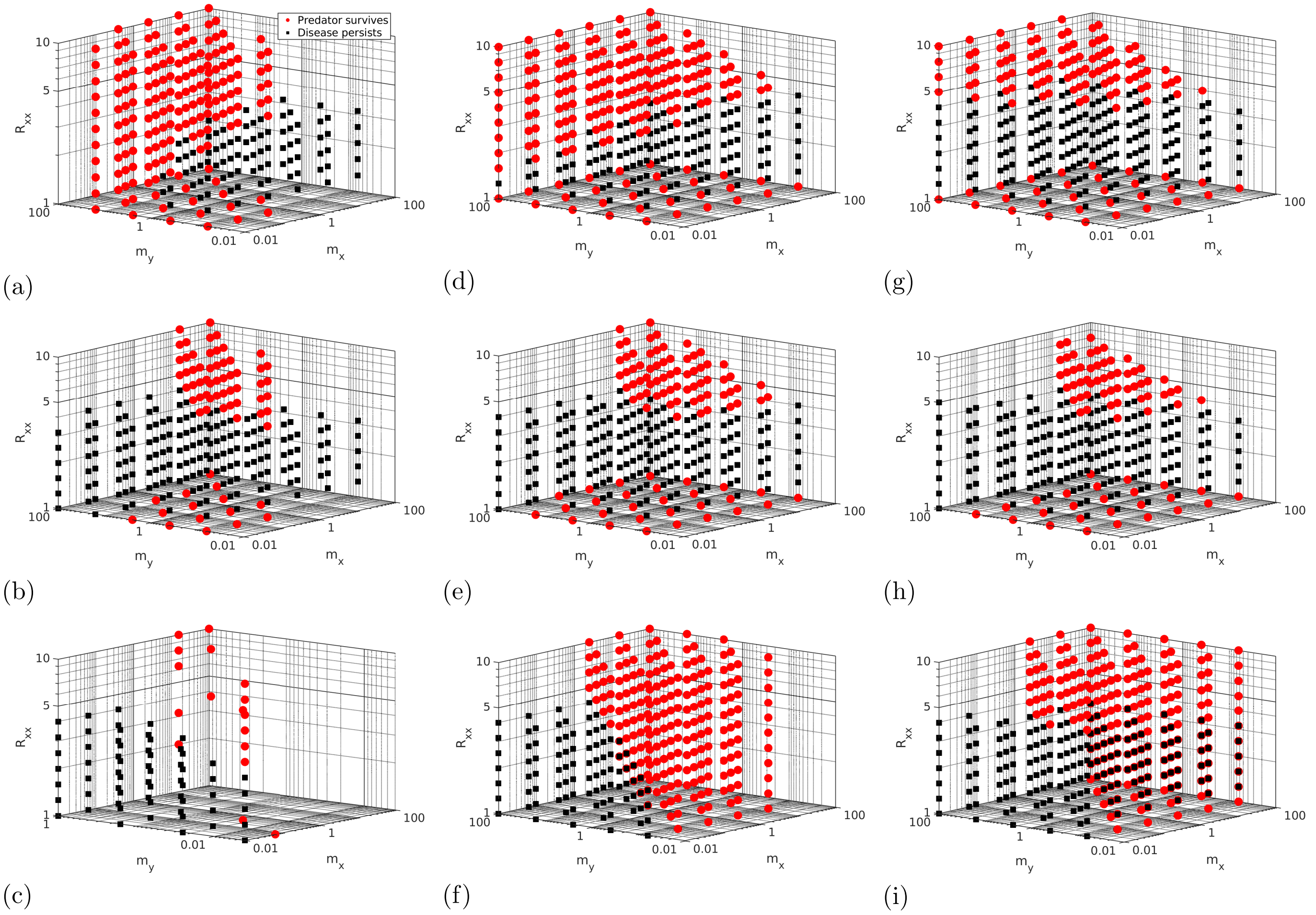


Figure 1: Plots showing the distribution of predator survival (red circles) and disease persistence (black squares) scenarios in parameter space for the three versions of our model. The coordinates of the markers indicate the parameter values leading to the given scenario, with prey mass $m_x$ and predator mass $m_y$ in kg. Row 1 (a,d,g) shows the model with immune predators, row 2 (b,e,h) susceptible predators, and row 3 (c,f,i) the model with two prey species, one immune and one susceptible, as well as susceptible predators. In the first column, figs. (a-c), we have set $\epsilon = 0.1K$. Here, the system appears to be unstable, at least when there are two prey species (fig. (c)). When $\epsilon = 0.3$ (figs. (d-f)), we get a distribution of predator survival and disease persistence similar to the one shown in the main text of this paper. This remains true for $\epsilon = 0.95K$ (figs. (g-i)). When $\epsilon > K$, the predator reproduction rate remains smaller than the death rate for all possible prey densities, and the model is therefore invalid for these $\epsilon$. We see that in the range from $\epsilon = 0.3K$ to $\epsilon \approx K$, the exact value of $\epsilon$ makes little difference, and we are therefore justified in taking it to be $K/2$.

### Outcomes for different prey-predator and predator-predator infectivities

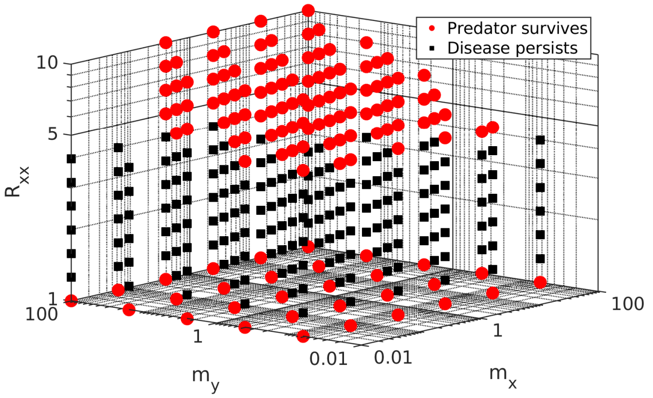


(a)

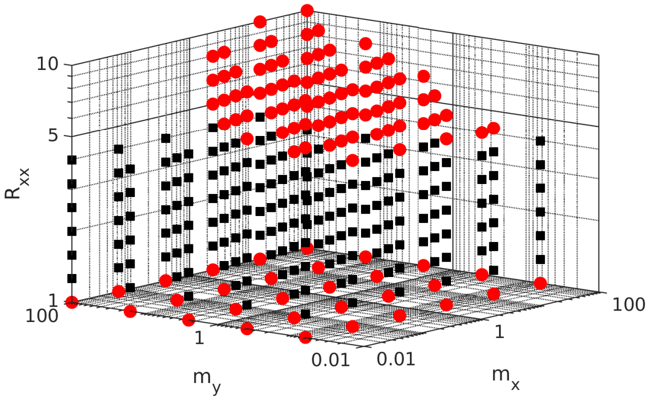


(c)

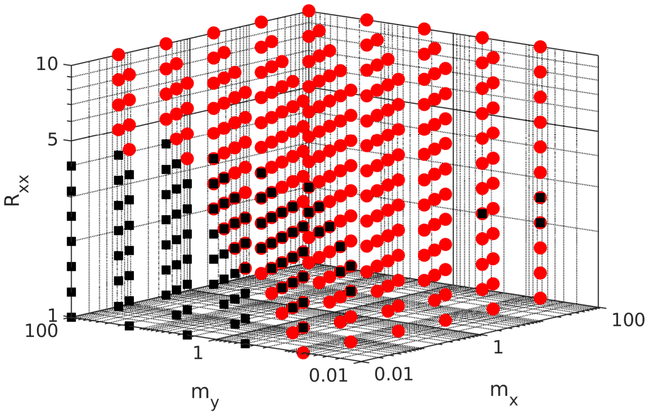


(b)

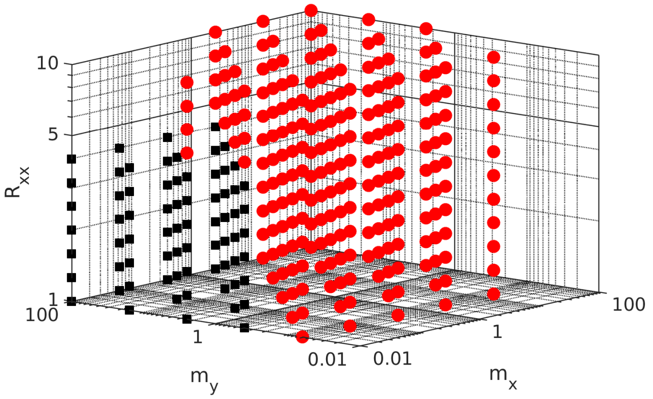


(d)

Figure 2: The effects of varying the infection probability when predators eat prey, $p_I$. The locations of the red dots in parameter space indicate parameter values leading to predator survival, while black squares indicate parameter values leading to disease persistence. (a) shows the one-prey system for $p_I = 0.1$ and (b) shows the two-prey system, also for $p_I = 0.1$. (c)-(d) show the same two systems for $p_I = 0.5$.

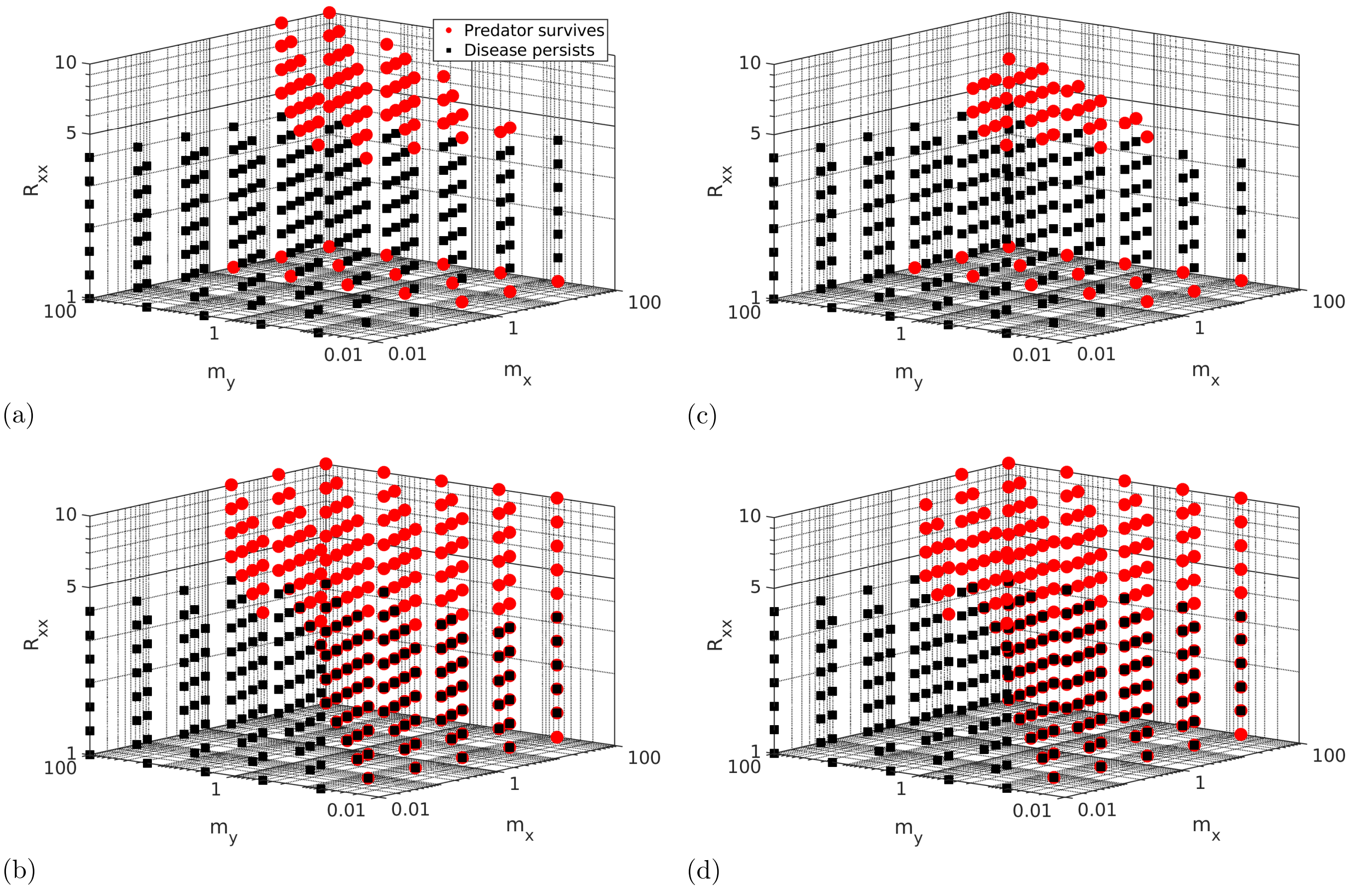


(a)

(c)

(b)

(d)

Figure 3: Plots demonstrating the effect of varying the rate of infection from predator to predator, here expressed via the parameter $R_{yy}$ which we have defined as $R_{yy} \equiv \beta_{yy} S_{y,0}/\gamma_y$, where $y_0 = \alpha\epsilon/\phi$ is the initial density of predators at the Lotka-Volterra equilibrium without disease. It is analogous to the basic reproduction number for prey-prey infections, $R_{xx}$. (a) and (b) show the parameter values leading to predator survival (red dots) and disease persistence (black squares) for $R_{yy} = 1.26$ in the one-prey and two-prey models respectively. The coordinates of the markers indicate the parameter values. (c) and (d) show the outcomes for the same two models, but for $R_{yy} = 5.01$. Given the magnitude of change of $R_{yy}$, the changes in predator survival are relatively minor.
